# Bipartite recruitment of PCH2 and COMET to the synaptonemal complex drives chromosome axis reconstruction leading to crossover restriction

**DOI:** 10.1101/2022.03.23.485444

**Authors:** Chao Yang, Kostika Sofroni, Yuki Hamamura, Bingyan Hu, Hasibe Tunçay Elbasi, Martina Balboni, Maren Heese, Arp Schnittger

**Affiliations:** National Key Laboratory of Crop Genetic Improvement, Hubei Hongshan Laboratory, Huazhong Agricultural University, Wuhan, 430070, China; Department of Developmental Biology, Institute of Plant Science and Microbiology, University of Hamburg, Hamburg, 22609, Germany; Department of Meiosis, Max Planck Institute for Multidisciplinary Sciences, 37077, Göttingen, Germany

## Abstract

Chromosome axis-associated HORMA domain proteins (HORMADs), e.g., ASY1 in Arabidopsis, are crucial for meiotic recombination. ASY1, as other HORMADs, is assembled on the axis at early meiosis and depleted when homologous chromosomes synapse. Puzzlingly, both processes are catalyzed by AAA+ ATPase PCH2 together with its cofactor COMET. Here, we show that the ASY1 remodeling complex is temporally and spatially differently assembled. While PCH2 and COMET appear to directly interact in the cytoplasm in early meiosis, PCH2 is recruited by the transverse filament protein ZYP1 and brought to the ASY1-bound COMET assuring the timely removal of ASY1 during chromosome synapsis. Since we found that the PCH2 homolog TRIP13 also binds to the ZYP1 homolog SYCP1 in mouse, we postulate that this mechanism is conserved among eukaryotes. Deleting the PCH2 binding site of ZYP1 led to a failure of ASY1 removal. Interestingly, the placement of one obligatory crossover per homologous chromosome pair, compromised by ZYP1 depletion, is largely restored in this engineered *zyp1* allele suggesting that crossover assurance is promoted by synapsis. In contrast, the engineered *zyp1* allele, similar to the *zyp1* null mutant, showed elevated type I crossover numbers indicating that PCH2-mediated eviction of ASY1 from the axis restricts crossover formation.

## Introduction

Sexual reproduction relies on the generation of gametes that contain only half of the genetic material of the parental cells; this reduction in chromosome number is achieved through meiosis. Besides its role in maintaining genome size over generations, meiosis generates genetic diversity through a random selection of either the maternal or the paternal homologous chromosome (homolog) and by an exchange of DNA segments between homologs by meiotic recombination.

Meiotic recombination is initiated by the formation of programmed double-strand breaks (DSBs) catalyzed by a complex containing the topoisomerase-like protein SPO11 and other accessory proteins (Wang & Copenhaver, 2018; Mercier *et al*, 2015; Vrielynck *et al*, 2021; Keeney, 2001). DSBs are resected by the MRN/MRX complexes, leading to single-strand DNA ends, which are bound by the recombinases RAD51 and DMC1. These recombinases promote the search for and initial invading steps into the sister chromatid of either the same chromosome or the homolog, resulting in either inter-sister or inter-homolog intermediates. The inter-homolog intermediates become associated with the ZMM proteins (including MSH4, MSH5, SHOC1(ZIP2), HEI10 (ZIP3), ZIP4, and MER3 in Arabidopsis) to form double-Holliday junctions (dHJs) that can be subsequently resolved either into class I (interference-sensitive) crossovers (COs) through the ZMM pathway together with two MutLg proteins, MLH1 and MLH3, or processed into non-crossovers (NCOs) by other mechanisms. In addition, a small fraction of inter-homolog intermediates is processed by another less-understood non-ZMM pathway (including the endonuclease MUS81) to form class II (interference-insensitive) COs (Mercier *et al*, 2015; Wang & Copenhaver, 2018; Hunter, 2015). As a consequence of COs, physical linkages (chiasmata) between homologs are formed at late prophase I that ensure the equal segregation of homologs during the first meiotic division (meiosis I) (Hunter, 2015; Wang & Copenhaver, 2018).

Recombination is critically dependent on the chromosome axis, a conserved meiosis-specific proteinaceous structure assembled along the entire length of chromosomes at early meiotic prophase I (Hunter, 2015; Wang & Copenhaver, 2018; Blat *et al*, 2002; Panizza *et al*, 2011; Zickler & Kleckner, 1999, 2015). The chromosome axis anchors and organizes the sister chromatids to form linear loop-arrays, enabling efficient DSB formation and inter-homolog recombination as observed in diverse taxa including, e.g., yeast, worms, mice and rice (Xue *et al*, 2019; Hollingsworth & Ponte, 1997; Niu *et al*, 2005; Goodyer *et al*, 2008; Carballo *et al*, 2008; Hunter, 2015).

The chromosome axis is composed of multiple proteins. In Arabidopsis, several axial components have been identified, including cohesin complexes, which build the basic scaffold of the axis; the HORMA domain-containing protein (HORMAD) ASY1 (homolog of Hop1 in yeast, HORMAD1/2 in mammals); and two coiled-coil proteins known as ‘axis core’, i.e., ASY3 (homolog of Red1 in yeast, SYCP2 in mammals) and ASY4 (homolog of SYCP3 in mammals) (Chambon *et al*, 2018; Armstrong, 2002; Wojtasz *et al*, 2009; Hollingsworth & Johnson, 1993; Fukuda *et al*, 2010; Lammers *et al*, 1994; Ferdous *et al*, 2012). Plants deficient in any of these components show strong defects in CO formation, highlighting the importance of the chromosome axis in recombination (Lambing *et al*, 2020b; Chambon *et al*, 2018; Sanchez-Moran *et al*, 2007; Ferdous *et al*, 2012; Armstrong, 2002).

The HORMADs play important roles in meiotic recombination in many organisms, including yeast, mammals, and plants (Niu *et al*, 2005; Carballo *et al*, 2008; Sanchez-Moran *et al*, 2007; Daniel *et al*, 2011; Wojtasz *et al*, 2009). During meiosis I, they are dynamically present on chromosomes in a stage-dependent manner, namely being assembled on chromosomes before synapsis and getting depleted from the axis when homologous chromosomes synapse (Yang *et al*, 2020b; Lambing *et al*, 2015; Chen *et al*, 2014; Roig *et al*, 2010).

Recently, it has been shown in Arabidopsis that PCH2 orchestrates both the chromosomal recruitment and dissociation of ASY1/Hop1 by mediating its conformational change (Balboni *et al*, 2020; Yang *et al*, 2020a; Raina & Vader, 2020). This mechanism is likely also present in mice and budding yeast. In Arabidopsis, PCH2 requires the cofactor COMET. Whether the necessity for a co-factor is conserved is currently not clear since no COMET homolog has so far been identified in budding yeast and a role of COMET in meiosis in mice has not been described so far.

While recently light has been shed on the role of the PCH2-COMET complex in ASY1 recruitment at early prophase I (Balboni *et al*, 2020; Yang *et al*, 2020a), the question of how this complex specifically dissociates ASY1 from the synapsed axes remains unclear. In particular, it is not understood how the two functions of PCH2 could be temporally separated, i.e., the assistance in ASY1’s recruitment to and the facilitation of the dissociation of ASY1 from the chromosome axis (Balboni *et al*, 2020; Yang *et al*, 2020a; Lambing *et al*, 2015).

Here, we demonstrate that nucleation of PCH2 on the synapsed axes is neither dependent on the chromosome axis components ASY1 and ASY3 nor on the PCH2 cofactor COMET but relies on the installation of ZYP1. We further show that PCH2 is recruited to the synapsed regions by directly binding to the C-terminal region of ZYP1, which is supposed to be proximal to the axis (Capilla-Pérez *et al*, 2021). Deletion of the PCH2-binding sequence in ZYP1 leads to a failure of PCH2’s nucleation on the synaptonemal complex (SC) and an extended presence of ASY1 on the chromosome axis. Furthermore, our results shed light on the hypothesized necessity of ASY1 removal from the axis for the complete polymerization of the SC and prevention of excess class I CO formation. Finally, we found that TRIP13, the ortholog of PCH2 in mouse, also directly binds to the C-terminus of the mouse ZYP1 homolog SYCP1, suggesting that the here-identified mechanism controlling ASY1 dynamics could also apply to other organisms, such as mammals.

## Materials and methods

### Plant materials

The Arabidopsis thaliana accession Columbia (Col-0) was used as the wild-type reference throughout this research. The T-DNA insertion lines SALK_046272 (*asy1-4*) (Yang *et al*, 2020b), SAIL_423H01 (*asy3-1*) (Ferdous *et al*, 2012), SALK_040213 (*zyp1a*) (Higgins *et al*, 2005), SALK_050581 (*zyp1b*)) (Higgins *et al*, 2005), *mlh1-3* (SK25975), and SALK_031449 (*pch2-2*) (Lambing *et al*, 2015) were obtained from the Salk Institute Genomics Analysis Laboratory (SIGnAL, http://signal.salk.edu/cgi-bin/tdnaexpress) via NASC (http://arabidopsis.info/). *The PRO_PCH2_:PCH2:GFP, PRO_COMET_:COMET:GFP, PRO_ASY3_:ASY3:RFP*, and *PRO_MLH1_-MLH1:GFP* reporter constructs were generated and described previously (Yang *et al*, 2020b; Balboni *et al*, 2020; Pochon *et al*, 2022). All plants were grown in growth chambers with a 16 h light/21°C and 8 h dark/18°C at the humidity of 60%.

### Plasmid construction and plant transformation

To generate the *PRO_ZYP1B_:ZYP1B^Δ661-728^* line, the coding sequence for the amino acids 661-725 in ZYP1B was deleted via PCR using the entry clone *PRO_ZYP1_:ZYP1B/pDONR221* previously generated as a template. The PCR fragments were ligated and subsequently integrated into the destination vector pGWB501 by gateway LR reaction. For creating the yeast two-hybrid construct of *ZYP1B-BD*, the full length CDS of ZYP1B was amplified via PCR using primers containing a *Nco*I (forward primer) and a *Bam*HI (reverse primer) restriction sites (Supplementary table 1). The PCR fragments were inserted into the *pGBKT7* vector via restriction-mediated cloning. To generated the ZYP1B truncations (*ZYP1B^1-300^-BD, ZYP1B^301-600^-BD, ZYP1B^601-856^-BD*), the coding sequence of relevant fragments was amplified via PCR using primers flanked with attB1 and attB2 sites followed by the gateway BP reaction with the *pDONR223* vector (Supplementary table 1). The other ZYP1B truncations (*ZYP1B^601-725^-BD, ZYP1B^726-834^-BD, ZYP1B^601-660^-BD*, and *ZYP1B^661-725^-BD*) were constructed by deleting the additional coding sequences of ZYP1B using the entry clones of *ZYP1B^601-856^-BD, ZYP1B^726-856^-BD*, or *ZYP1B^601-725^-BD* as the PCR template (Supplementary table 1). Next, the resulting entry clones were integrated into the *pGBKT7-GW* vector by a gateway LR reaction. The PCH2-related AD (activation domain) constructs were described previously (Balboni *et al*, 2020).

### Yeast two-hybrid assay

To perform the yeast two-hybrid interaction test, the relevant AD and BD constructs were co-transformed into the auxotrophic yeast strain AH109 using the polyethylene glycol/lithium acetate method according to the manufacture’s manual (Clontech). Yeast cells haboring the relevant combinations of constructs were dotted on plates with double (-Leu-Trp), triple (-Leu-Trp-His) and quadruple (-Leu-Trp-His-Ade) synthetic dropout medium to assay growth.

### Cytological analysis

The imaging of reporters in male meiocytes was performed according to (Prusicki *et al*, 2019). In brief, Arabidopsis anthers harboring the fluorescent reporters at appropriate meiotic stage were dissected and immediately imaged using a Leica TCS SP8 inverted confocal microscope. The meiotic stages were determined by the criteria including chromosome morphology, nucleolus position, and cell shape (Prusicki *et al*, 2019).

Meiotic chromosome spread analysis was performed according to (Yang *et al*, 2020a). In brief, fresh flower buds were fixed in Carnoy’s fixative (ethanol: acetic acid = 3: 1) for 48 h at 4°C followed by two times of washing with 75% ethanol, and then stored in 75% ethanol at 4°C. For chromosome spreading, flower buds at appropriate stage were initially digested in the enzyme solution (10 mM citrate buffer containing 1.5% cellulose, 1.5% pectolyase and 1.5% cytohelicase) for 3 h at 37°C. Next, single flowers were dissected and smashed with a bended needle on the microscopy slide (note: avoid drying the samples). The spreading step was performed on a 46°C hotplate with 10 μl of 45% acetic acid and then the slide was rinsed with ice-cold Carnoy’s fixative. After drying at 37°C for 12h, the slides were mounted with anti-fade DAPI solution (Vector Laboratories).

Immunostaining experiment was performed as described previously (Yang *et al*, 2020a). Briefly, fresh flower buds at appropriate meiotic stage were dissected and macerated in 10 μl digestion enzyme mixture (0.4% cytohelicase, 1% polyvinylpyrrolidone, and 1.5% sucrose) on the poly-lysine coated slide for 7 mins in a moisture chamber at 37°C followed by a squashing. Next, the samples were incubated further for 8 mins after adding another 10 μl enzyme solution. Subsequently, the samples were smashed thoroughly in 20 μl 1% Lipsol solution. Next, the spreading was performed after adding 35 μl fixative (4% (w/v) paraformaldehyde, pH 8.0) and the slides were dried at room temperature for 2–3h. For the immunostaining, the slides were washed three times in PBST buffer and blocked in PBST containing 1% BSA for 1 h at room temperature in a moisture chamber. Next, the slides were incubated with relevant antibodies (anti-GFP: gb2AF488 from Chromotek (1:300 dilution), anti-ASY1 (1:500 dilution), anti-ZYP1 (1:500 dilution) at 4°C for 48 h. After three times of washing (10 mins each) in PBST, the slides were incubated with fluorescein-conjugated secondary antibodies for 24h at 4°C. Next, the DNA was counterstained with antifade DAPI solution (Vector Laboratories) following three times of washing. Images were acquired using the Leica TCS SP8 inverted confocal microscopy.

## Results

### Nucleation of PCH2 on the SC is independent of ASY1, ASY3 and COMET

In the wildtype, PCH2 is recruited to the SC coinciding with chromosome synapsis at zygotene and stays along synapsed chromosomes throughout pachytene (Figure 1A, B) (Yang *et al*, 2020b; Lambing *et al*, 2015). We therefore asked whether this localization pattern depends on the chromosome axis protein ASY1 and/or ASY3 as parts of the lateral element (LE) of the SC. To address this question, we introduced a previously generated functional *PCH2* reporter line, *PCH2:GFP* (Yang *et al*, 2020b), into *asy1* and *asy3* mutants and followed the localization of PCH2 in male meiocytes by laser scanning confocal microscopy. In *asy1* and *asy3* mutants, chromosome synapsis and formation of the SC are largely, yet not completely, impaired (Lambing *et al*, 2020a; Ferdous *et al*, 2012). Since we could detect short stretches of PCH2:GFP (Figure 1B), we conclude that PCH2 can still be recruited to the remnants of the SC present in these mutants, consistent with a previous report using immuno-detection techniques (Lambing *et al*, 2015). Thus, ASY1 and ASY3 are not required for the localization of PCH2 to the SC.

**Figure 1.**
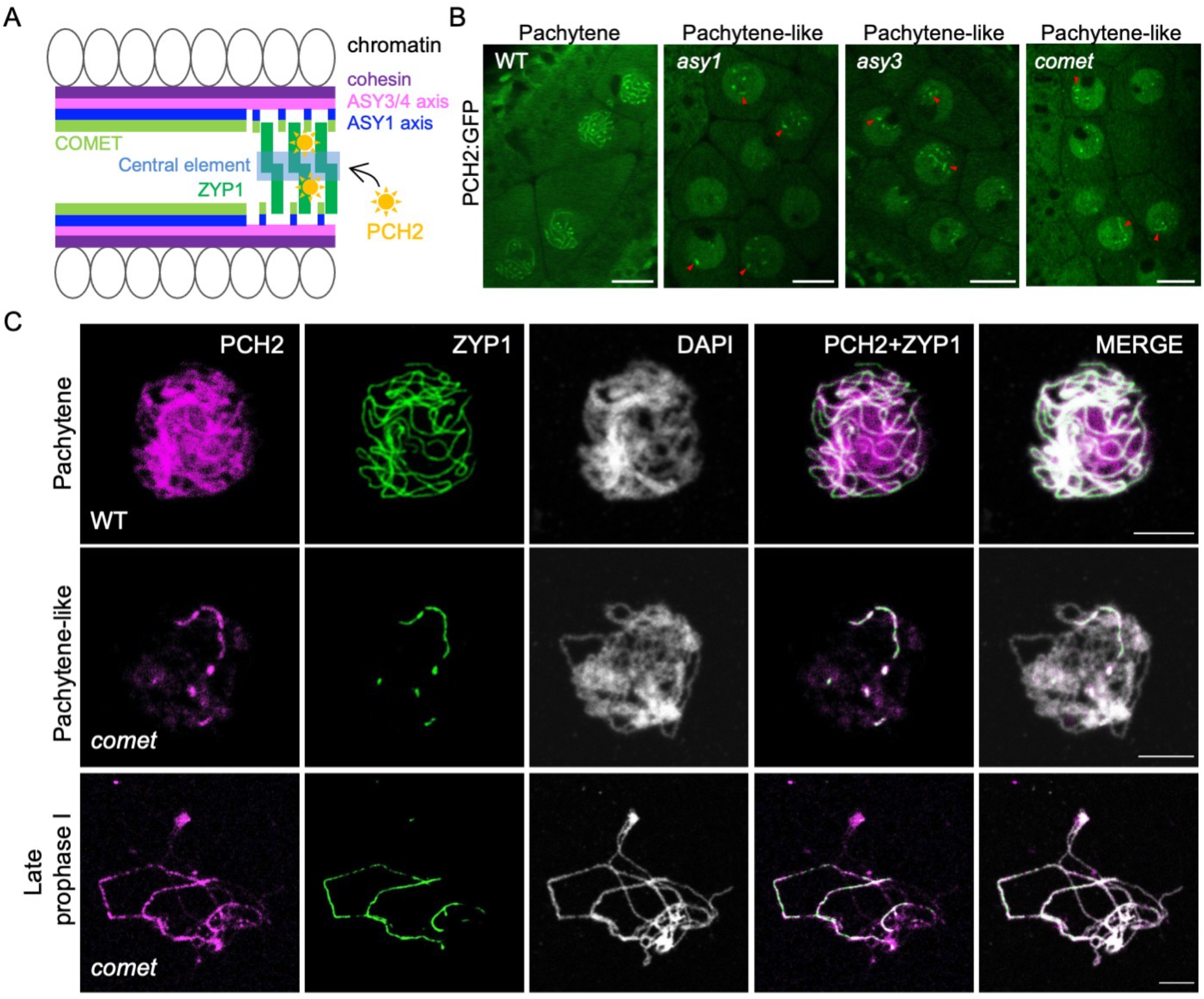
Recruitment of PCH2 to the synaptonemal complex is independent of the chromosome axis and its co-factor COMET. (A) A schematic representation of the structure of synapsing chromosomes and the key proteins involved. (B) Localization of PCH2:GFP in the male meiocytes of wild-type (WT), *asy1, asy3*, and *comet* mutant plants at pachytene or pachytene-like stages using confocal laser scanning microscope. Red arrowheads indicate the short stretches of PCH2 signal localizing at synapsed chromosomal regions. Bars: 10 μm. (C) Co-immunolocalization of PCH2 and ZYP1 in WT and *comet* mutants at pachytene or pachytene-like stages and at late prophase I. Bars: 5 μm.

Previously, we found that COMET, in its function as an adaptor for PCH2, is associated with chromosomes by binding likely directly to ASY1 in early prophase (Balboni *et al*, 2020). Therefore, we wondered if the recruitment of PCH2 to the SC relies on COMET. In *comet* mutants, chromosome synapsis is also largely compromised, but occasional stretches of SC can be formed (Balboni *et al*, 2020). Similar to the situation in *asy1* and *asy3* mutants, we found that PCH2 could still assemble into short thread-like structures in male meiocytes of *comet* mutants (Figure 1B). We confirmed this finding by coimmunolocalization of PCH2 and ZYP1 in *comet* mutants (Figure 1C).

Taken together, these results suggest that PCH2’s nucleation on the SC is not dependent on the axis-associated proteins ASY1, ASY3, and COMET, yet tightly correlates with chromosome synapsis and SC formation.

### PCH2 does not nucleate on paired chromosomes in the absence of ZYP1

Next, we asked whether PCH2’s nucleation on the SC might rely on ZYP1, which is up to now the only component identified in plants belonging to the central region of the SC. However, ZYP1 function in Arabidopsis is distributed to two redundantly-acting and in tandem arranged genes, *ZYP1A* and *ZYP1B*, which are positioned tail to tail with less than 2 kb in total. Since no *zyp1a zyp1b* double mutant existed when we started this work, we mutated *ZYP1B* via CRISPR-Cas9 in the background of a *zyp1a* T-DNA insertion line (SALK_040213), leading to four different double mutant combinations: *zyp1a/b-1, zyp1a/b-2, zyp1a/b-3*, and *zyp1a/b-4,* harboring next to the T-DNA insertion in *ZYP1A* a C insertion, a 17 bp deletion, an A insertion, and a TG deletion in *ZYP1B*, respectively (Supplemental figure 1A). All these mutations in *ZYP1B* lead to shifts in the reading frame and the formation of premature stop codons, thus likely representing null mutants (Supplemental figure 1A). Consistent with a complete loss of ZYP1 function in these alleles, we did not detect a ZYP1 signal in any of the four *zyp1a/b* double mutant combinations by immunodetection of ZYP1 using an antibody that recognizes both paralogs (Supplemental figure 1B).

Next, we performed a detailed phenotypic analysis of the *zyp1a/b* mutants. However, while this work was in progress, two other reports presented the analysis of *zyp1a/b* double mutants (France *et al*, 2021; Capilla-Pérez *et al*, 2021). Our results of the fertility and chromosome behavior in *zyp1a/b* mutants shown in Supplemental figure 1C-G and 2A are largely consistent with these two reports except for reduced pollen viability, which we detected in our assays. Briefly, we found that, compared to the wildtype, there were no obvious growth defects in *zyp1a/b* double mutants. Additionally, no obvious difference in the silique length of the double mutants in comparison with the wildtype could be seen, suggesting that *zyp11a/b* mutants are largely fertile (Supplemental figure 1C). However, *zyp1a/b* mutants showed a slight reduction in pollen viability amounting to ~2-3% in comparison to the wildtype and *zyp1a* single mutants (one-way ANOVA with Dunnett correction for multiple comparison, P < 0.001), (Supplemental figure 1D and E. In addition, we observed a significant though mild reduction in seed set (Supplemental figure 2F and G) (P < 0.001), suggesting very likely slight defects in female meiosis and/or abortion of early embryos.

Consistent with the previous reports (France *et al*, 2021; Capilla-Pérez *et al*, 2021), chromosome spread analyses revealed that chromosome pairing and coalignment can function largely independently of ZYP1 in Arabidopsis (Supplemental figure 2A[a-l]). In contrast, ZYP1 is required for tight homolog interaction and CO assurance as judged by the presence of univalents in *zyp1a/b* mutants (17.6% in *zyp1a/b-1*, n = 74 meiocytes; 10.3% in *zyp1a/b-2,* n = 78 meiocytes; 12.5% in *zyp1a/b-3*, n = 64 meiocytes; 14.3% in *zyp1a/b-4*, n = 21 meiocytes) (Supplemental figure 2A[m-s] and B).

The finding that loss of *ZYP1* does not affect the pairing/coalignment of chromosomes allowed us to assess whether pairing as opposed to the presence of ZYP1 is required for the recruitment of PCH2 to chromosomes. To this end, we introduced the *PCH2:GFP* reporter, along with an *ASY3:RFP* construct to stage meiotic nuclei (Yang *et al*, 2019), into the *zyp1a/b-1* double mutants. While PCH2 nucleated on synapsed chromosomes at pachytene in the wildtype, we observed only a diffuse GFP signal in the *zyp1a/b-1* mutant. ASY3:RFP, on the other hand, showed a wild-type-like localization pattern on paired chromosomes (Figure 2A, B, Supplemental figure 4A).

**Figure 2.**
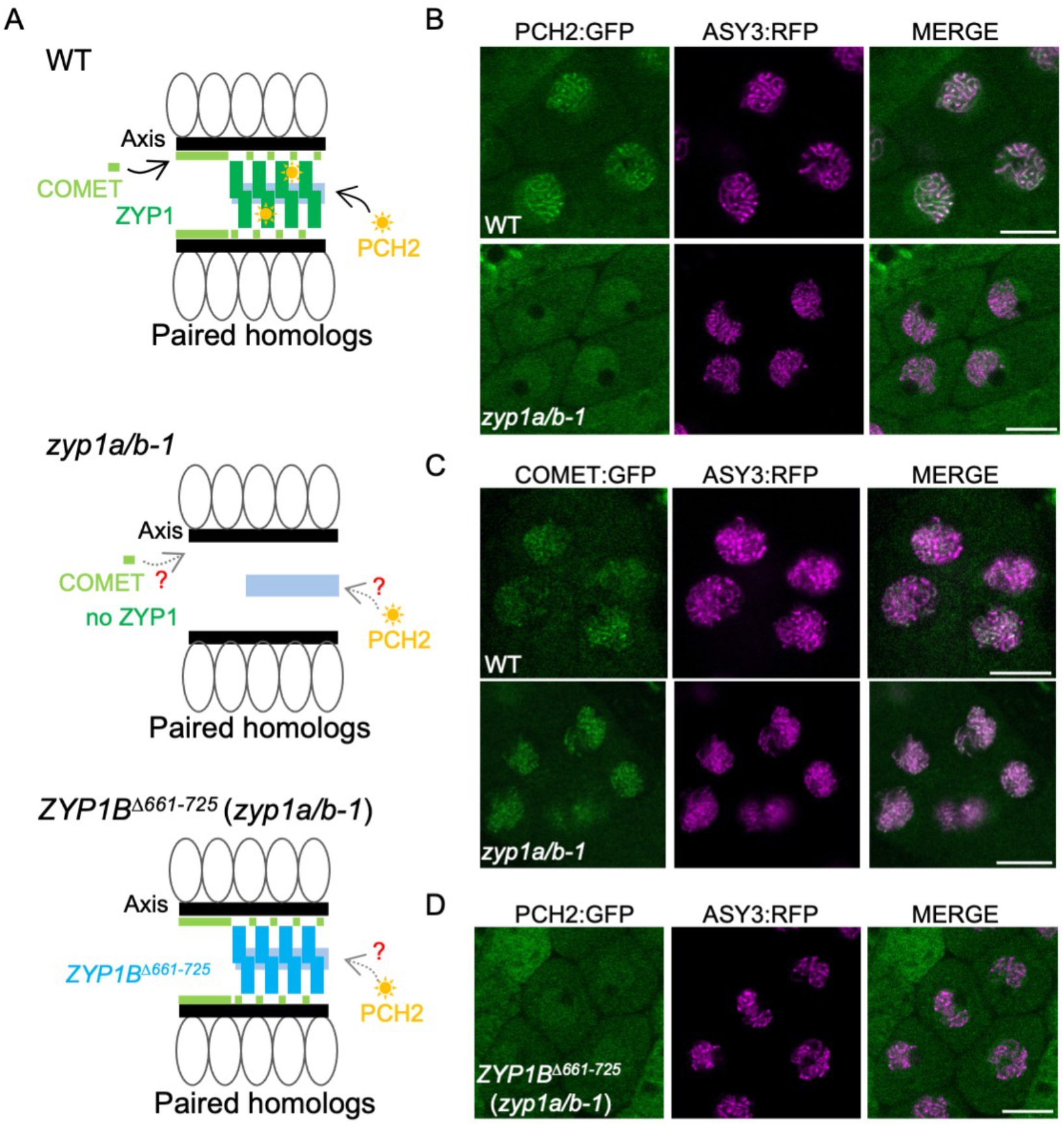
Recruitment of PCH2 to the synaptonemal complex is dependent on ZYP1. (A) Schematic depiction of the structure of synapsing/co-aligning chromosomes in WT, *zyp1a/b-1*, and *ZYP1B*^Δ*661-725*^ (*zyp1a/b-1*) mutant plants. (B) Localization patterns of PCH2:GFP together with ASY3:RFP in male meiocytes of WT and *zyp1a/b-1* mutant plants at pachytene or pachytene-like stages. Bars: 10 μm. (C) Localization patterns of COMET:GFP together with ASY3:RFP in male meiocytes of WT and *zyp1a/b-1* mutant plants at early prophase. Bars: 10 μm. (D) Localization pattern of PCH2:GFP together with ASY3:RFP in male meiocytes of WT and *ZYP1B*^Δ*661-725*^ (*zyp1a/b-1*) mutant plants at pachytene-like stage. Bars: 10 μm.

Since COMET functions as the cofactor of PCH2 in Arabidopsis (Balboni *et al*, 2020), we also asked whether the absence of ZYP1 would influence the chromosome association of COMET, too. To address this, a previously generated functional *COMET* reporter, *COMET:GFP* (Balboni *et al*, 2020), was introduced into the *zyp1a/b-1* mutants together with *ASY3:RFP.* Following the localization of COMET in the male meiocytes of wild-type and *zyp1a/b-1* mutant plants revealed that COMET’s association with chromosomes was not affected in *zyp1a/b-1* mutants (Figure 2C).

Overall, these data show that the two components of the ASY1 dissociation machinery are brought together via two different pathways: COMET’s localization is independent of the TF protein ZYP1 but relies on the LE and ASY1 itself (Balboni *et al*, 2020); conversely, PCH2 relies on ZYP1 for SC localization but does neither require ASY1 nor ASY3 to be present on the axis.

### PCH2 directly interacts with the C-terminus of ZYP1B

The findings above raised the question whether PCH2 would directly bind to ZYP1. To answer this question, we performed yeast two-hybrid assays. An initial assay between the full length PCH2 protein and the entire ZYP1B protein, as a representative of the two ZYP1 proteins, did not reveal an interaction (Figure 3A, B). However, the binding of two truly interacting proteins *in vivo* can be masked when tested *in vitro* or in a heterologous system due to incorrect folding, improper conformation, or lack of secondary modifications as, for instance, seen for the interaction between ASY1 and ASY3 (Yang *et al*, 2020b). Therefore, we first tested the binding of ZYP1B to two earlier generated PCH2 truncations: PCH2^1-130^, which we have previously found to interact with COMET, and PCH2^131-467^, which does not bind to COMET (Balboni *et al*, 2020). Nevertheless, again no interaction could be detected (Figure 3B). However, when we next divided ZYP1B into three parts (1-300 aa, 301-600 aa, and 601-856 aa), we found that the C-terminal fragment (ZYP1B^601-856^) interacted with PCH2^1-130^ (Figure 3B).

**Figure 3.**
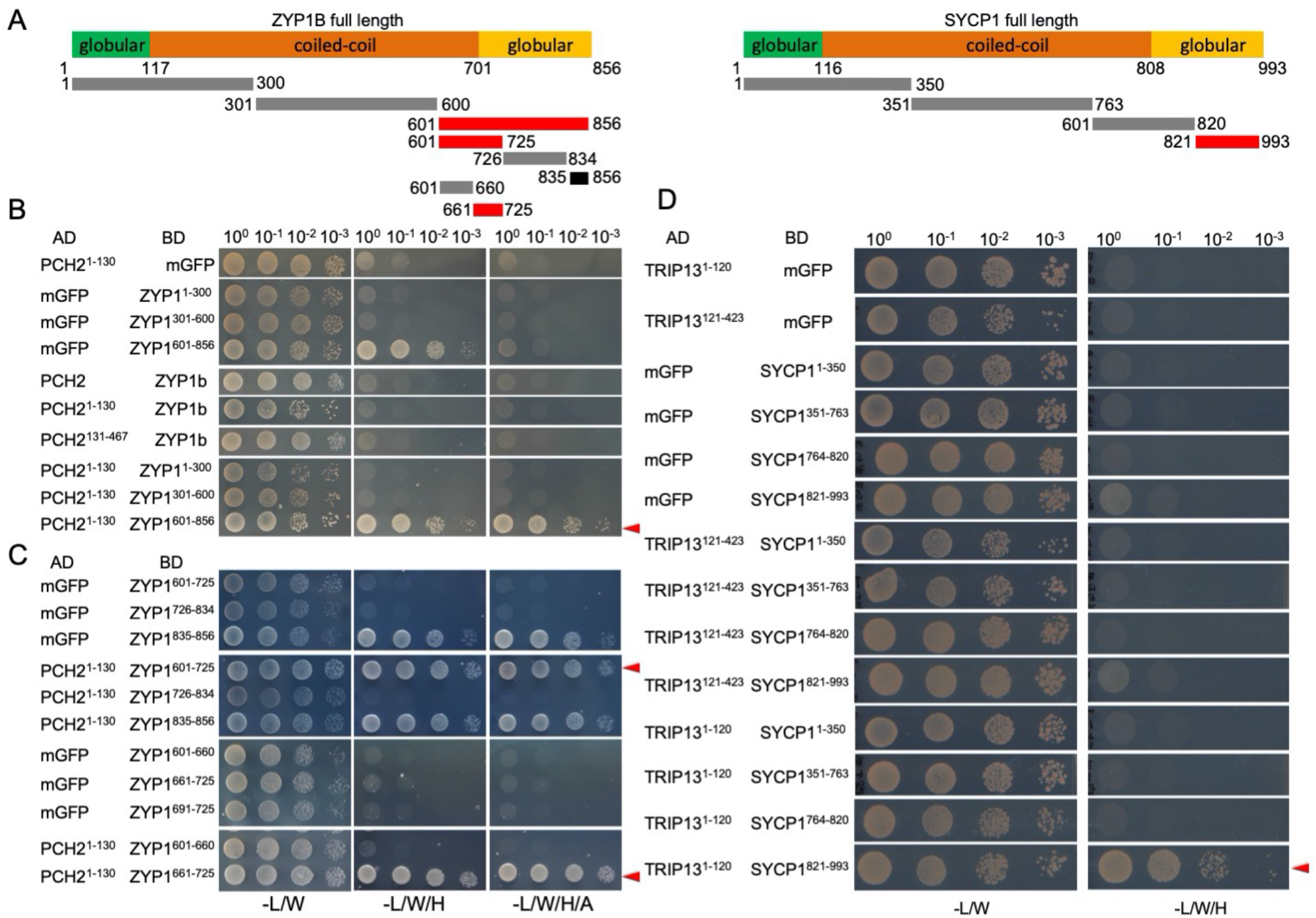
PCH2/TRIP13 interacts with ZYP1/SYCP1. (A) Schematic graph of Arabidopsis ZYP1B and mouse SYCP1 showing the N- and C-terminal globular domains and the central coiled-coil domains. The lines below indicate the truncations used in the yeast two-hybrid interaction assay. Red lines indicate the positive interaction and black line indicates the autoactivation. (B-D) Yeast two-hybrid assay of PCH2 and ZYP1 (B and C) or TRIP13 and SYCP1 (D) with different truncations. Red arrowheads indicate positive interactions.

To narrow down the binding domain of ZYP1B with PCH2, we further divided the C-terminal part of ZYP1B into three segments: 601-725 aa, 725-834 aa, and 835-856 aa. ZYP1B^835-856^ showed a strong autoactivation in our assay and hence, could not be evaluated. While PCH2^1-130^ did not bind to ZYP1B^726-834^, a strong interaction of ZYP1B^601-725^ with PCH2^1-130^ was observed (Figure 3C). It is worth mentioning that ZYP1B^661-725^ is almost identical to ZYP1A in the region between amino acids 661 and 725 (Supplemental figure 3), with only one differing amino acid. Thus, we conclude that PCH2 binds with its N-terminal region (1-130) to the C-terminal parts of ZYP1B (661-725) and likely ZYP1A (661-725).

### A separation-of-function mutant of *ZYP1B* reveals the necessity of the SC-localized PCH2 for ASY1 depletion

Seeing the elaborated system that assures the tightly temporally controlled presence of ASY1 at the chromosome axis, we next asked what the consequences of an alteration of its residence time are. The failure of bringing the ASY1 dissociation complex together to the paired/coaligned chromosomes explains the previous findings that loss of ZYP1 results in a prolonged/continuous presence of ASY1 on the axis (France *et al*, 2021; Capilla-Pérez *et al*, 2021). The entry of ASY1 into the nucleus followed by the formation of a linear ASY1 signal along chromosome axes in *zyp1a/b* mutants, reflecting the situation in the wildtype, suggests that the cytoplasm- and nucleoplasm-localized PCH2 is sufficient for nuclear targeting and full chromosome assembly of ASY1 (Supplemental figure 5). In accordance with the reports from Capilla-Pérez *et al* (2021) and France *et al* (2021), when meiocytes reached the pachytene-like stage (visualized by the DAPI-stained thick thread-like chromosomes), characterized by paired and coaligned homologous chromosomes, in *zyp1a/b* mutants we observed no obvious removal of ASY1 from the paired axes as revealed by immunodetection (Figure 4C and Supplemental figure 4B). This observation was confirmed by imaging the live male meiocytes of *zyp1a/b-1* mutants harboring a previously generated ASY1:GFP reporter together with the above-used ASY3:RFP (Yang *et al*, 2020b) (Supplemental figure 5). This gave rise to the hypothesis that the recombination defects of *zyp1* mutants are at least partially due to the prolonged presence of ASY1 on the axis.

**Figure 4.**
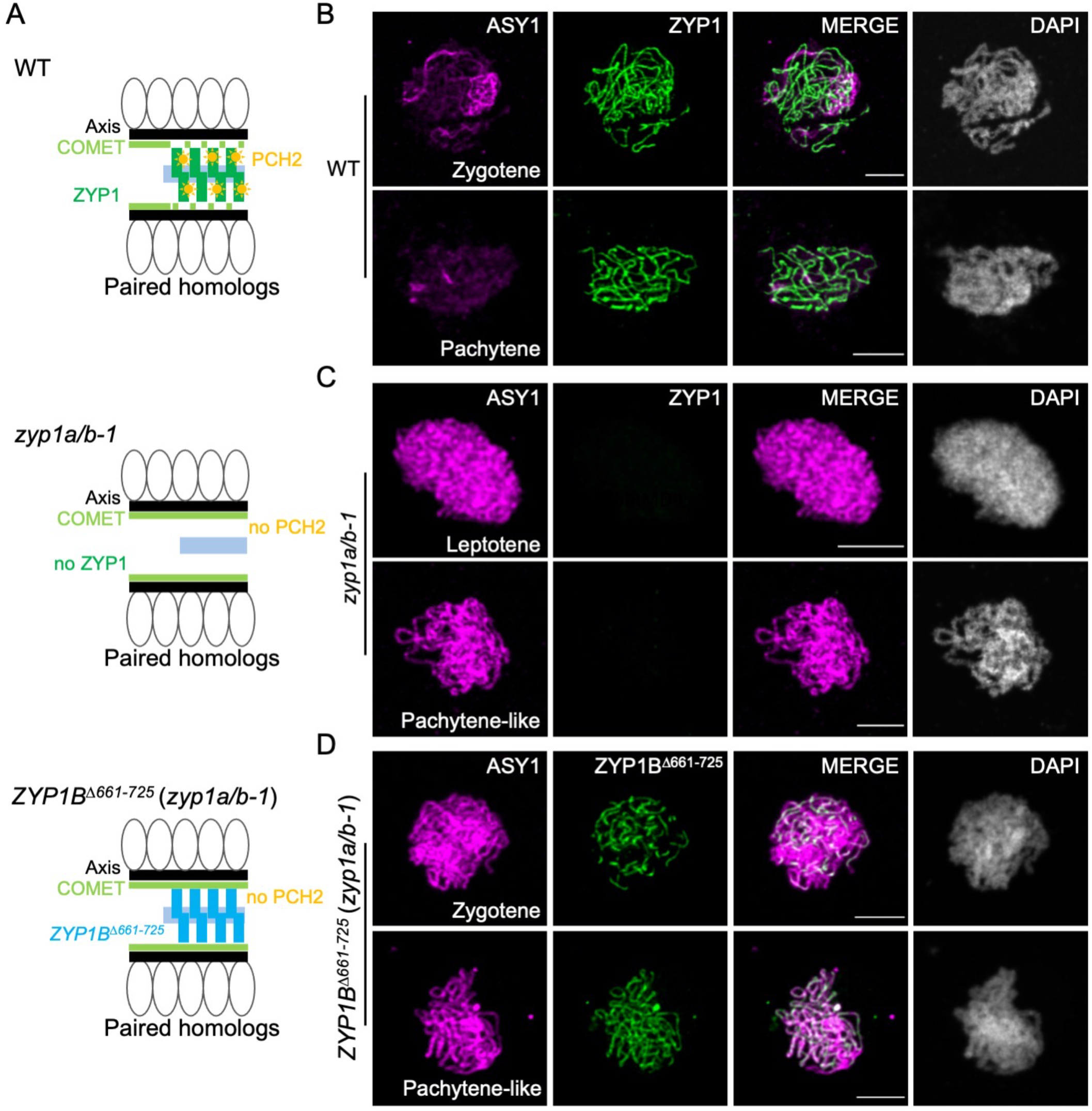
Absence of ZYP1 results in the deficient removal of ASY1 from the chromosome axis. (A) Schematic depiction of the structure of paired chromosomes in WT, *zyp1a/b-1*, and *ZYP1B*^Δ*661-725*^ (*zyp1a/b-1*) mutant plants. (B-D) Co-immunolocalization of ASY1 with ZYP1 or ZYP1B^Δ661-725^ in WT (B), *zyp1a/b-1* (C), and *ZYP1B^Δ661-725^* (*zyp1a/b-1*) (D) mutant plants at different prophase I stages. Bars: 5 μm.

However, the absence of ZYP1 and, with that, the lack of synapsis might have further and possibly indirect and not yet understood consequences, resulting in the observed increase in CO formation and independently of an altered ASY1 pattern (Capilla-Pérez *et al*, 2021; France *et al*, 2021). To tackle this problem, we generated a separation-of-function version of *ZYP1B* which keeps the ability to polymerize along the paired axes but cannot bind to PCH2. To this end, we used a previously generated *ZYP1B* genomic construct (Yang *et al*, 2019) and deleted the PCH2 binding domain in ZYP1B located between amino acids 661-725 (*PRO_ZYP1B_:ZYP1b^Δ661-725^*, called *ZYP1B^Δ661-725^*), as mapped above by the yeast two-hybrid assays (Figure 3C).

We first introduced the *ZYP1B*^*Δ661*-725^ construct into *zyp1a/b-1* mutants, called *ZYP1B*^Δ*661-725*^ (*zyp1a/b-1*) (Figure 2A), and found that the deletion of 661-725 aa still allowed the polymerization of ZYP1B on the SC as revealed by immunodetection of ZYP1 (Supplemental figure 6A). Despite the formation of a SC, PCH2 was not recruited to the chromosomes in *ZYP1B*^Δ*661-725*^ (*zyp1a/b-1*) at pachytene-like stages as determined by the chromosome morphology and cell shape (Figure 2D). Instead, PCH2: GFP showed a diffuse localization pattern in the nucleus in contrast to the wildtype (Figure 2B and D), corroborating that the domain between 661-725 in ZYP1 binds to PCH2 *in planta*. Importantly, we found by immunodetection that ASY1 could not be removed from the synapsed axes of *ZYP1BD*^Δ*661-725*^ (*zyp1a/b-1*) plants, resembling the situation in *zyp1a/b-1* double mutants (Figure 4D). This result was confirmed by the imaging of an ASY1:GFP reporter in male meiocytes of *ZYP1B*^*Δ661-725*^ (*zyp1a/b-1*) plants (Supplemental figure 6B). These data suggest that the ZYP1-dependent nucleation of PCH2 on the SC is crucial for ASY1 removal from the synapsing axes.

A detailed analysis of the SC in male meiocytes of ZYP1B^Δ661-725^ (*zyp1a/b-1*) plants by ZYP1 immunolocalization revealed that ZYP1 polymerization was nearly completed in only 3 out of 30 meiocytes. In the remaining 27 meiocytes, ZYP1 did not accumulate along the entire length of the chromosome axes throughout pachytene stage (Figure 5A). In contrast, the SC appeared to be completely assembled along with the regular depletion of ASY1 in wild-type plants expressing the *ZYP1B*^*Δ661-725*^ construct (n = 20 meiocytes) at pachytene stage (Figure 5B). These results suggest that the assembly of ZYP1B^Δ661-725^ *per se* on paired chromosomes has probably no dominant impact on the SC extension and that the incomplete assembly of ZYP1B^Δ661-725^ in *zyp1a/b-1* mutants is conceivably due to the compromised ASY1 removal from the paired axes. Thus, it is tempting to speculate that the extended presence of ASY1 on the axes interferes with SC formation.

**Figure 5.**
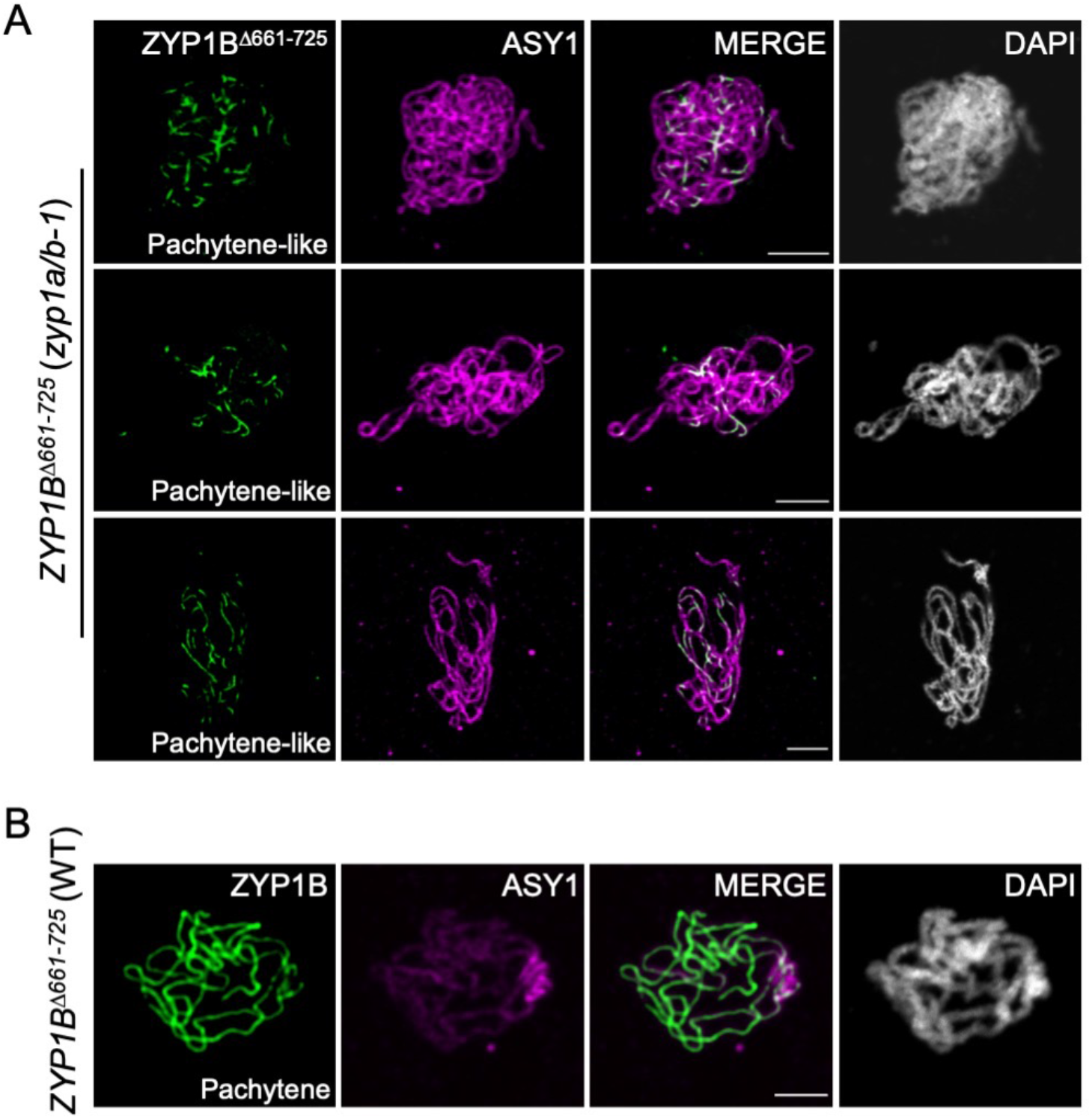
ZYP1^Δ661-725^ does not completely polymerize in *zyp1* mutants. (A) Co-immunolocalization of ZYP1B^Δ661-725^ and ASY1 in *ZYP1B*^Δ*661-725*^ (*zyp1a/b-1*) mutant plants at pachytene-like stage. Bars: 5 μm. (B) Co-immunolocalization of ZYP 1B^Δ661-725^ and ASY1 in WT plants at pachytene. Bars: 5 μm.

### *ZYP1b*^*Δ661*-725^ partially restores bivalent formation, yet still leads to an increase of type I COs

This *zyp1* separation-of-function allele allowed us then to evaluate the effects triggered by the complete loss of ZYP1 versus the prolonged axial presence of ASY1 in plants that form a SC. First, we compared the number of chiasmata in *zyp1a/b* mutants versus the *ZYP1B^Δ661-725^* (*zyp1a/b-1*) plants according to the cytological configuration of the bivalents at metaphase I in male meiocytes as previously described in (Santos *et al*, 2003). In the wildtype, the estimated minimum number of crossovers was 8.30 ± 0.89 (n = 50 meiocytes) per meiosis. This number was slightly, yet significantly (one way ANOVA with Tukey’s multiple comparison test) reduced to 7.76 ± 1.03 (n = 51meiocytes, p = 0.0319), 7.45 ± 0.92 (n = 38 meiocytes, p = 0.0004), and 7.52 ± 1.02 (n = 64 meiocytes, p = 0.0002) in *zyp1a/b-1, zyp1a/b-2, and zyp1a/b-3* mutants, respectively (Figure 6A). These data are in accordance with previously published results (Capilla-Pérez *et al*, 2021; France *et al*, 2021). In comparison, we found that the estimated minimum number of chiasmata in *ZYP1B^Δ661-725^* (*zyp1a/b-1*) plants (8.45 ± 0.97, n = 95 meiocytes) was slightly increased and reached wild-type levels. Interestingly, the number of meiocytes containing univalents significantly decreased from over 10% in *zyp1* double mutants (see above) to 4.2% (n = 120 meiocytes, P = 0.047, chi-square test) in *zyp1* mutants expressing *ZYP1b^Δ661-725^*, indicating that either ZYP1 or synapsis *per se* are able to promote the formation of the obligatory CO without ASY1 depletion

**Figure 6.**
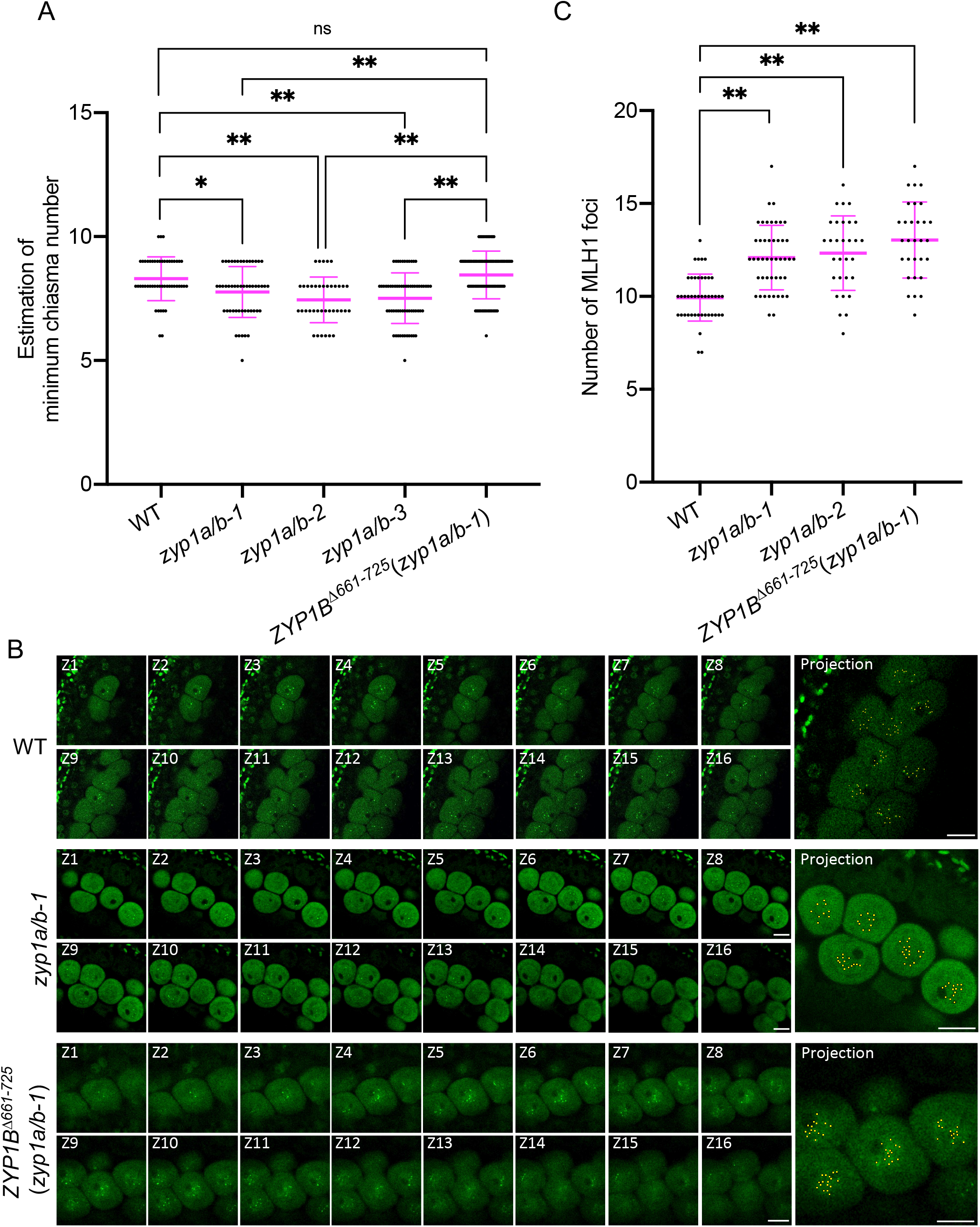
ZYP1 and ZYP1-mediated ASY1 removal regulate the formation of wild-type level of crossovers. (A) Scatter dot plot of the estimation of minimum chiasma number in WT, *zyp1a/b-1, zyp1a/b-2, zyp1a/b-3*, and *ZYP1B*^*Δ661-725*^ (*zyp1a/b-1*) mutant plants. (B) Z-stack imaging of MLH1:GFP in WT, *zyp1a/b-1*, and *ZYP1B*^*Δ661*-725^ (*zyp1a/b-1*) mutant plants at late prophase of male meiocytes. Sixteen z-stack images (Z1-Z16) with 0.8 μm interval are shown. All the MLH1 foci are indicated in the projection image with yellow dots. Bars: 10 μm. (C) Scatter dot plot of the number of MLH1 foci in WT, *zyp1a/b-1, zyp1a/b-2*, and *ZYP1B^Δ661-725^*(*zyp1a/b-1*) mutant plants. * and ** indicate significant differences at P < 0.05 and P < 0.01, respectively.

Previous reports showed that the number of type I COs increases in the absence of ZYP1, despite a reduced number of distinguishable chiasmata. This discrepancy can be explained by more closely located COs, as evidenced by sequencing, that cannot be resolved cytologically, for instance in *asy1* mutants (France *et al*, 2021; Capilla-Pérez *et al*, 2021) To understand whether the elevation of type I COs in *zyp1* mutants is due to the absence of ZYP1 *per se* or related to the defective ASY1 removal, we analyzed whether the ZYP1B^Δ661-725^-mediated formation of the SC, which fails to recruit PCH2 for ASY1 removal, would affect the number of type I COs in *zyp1* mutants. To answer this, we decided to count the number of MLH1 foci present in late pachytene cells. MLH1 is one of the MutLγ endonucleases crucial for the resolution of crossover intermediates and has been widely used in different organisms, including Arabidopsis, to estimate the number of type I crossovers. To achieve this, we employed a previously generated functional MLH1 reporter line (Pochon *et al*, 2022) and introgressed it into *zyp1a/b*, and *ZYP1BD^661-725^ (zyp1a/b-1)* mutant plants for MLH1 foci counting. To detect all of the MLH1 foci and properly count them, z-stack images were acquired with 0.8 μm interval (Supplemental movie 1 and 2). We found 9.93 ± 1.26 (n = 44 meiocytes) MLH1 foci per meiocyte in the wildtype (Figure 6B and C, Supplemental movie 1), consistent with previous reports using an immunodetection method (Chelysheva *et al*, 2010; Lambing *et al*, 2015; Ferdous *et al*, 2012). Similar to the findings in (France *et al*, 2021; Capilla-Pérez *et al*, 2021), this number slightly, yet significantly increased to 12.02 ± 1.79 (n = 46 meiocytes) and 12.33 ± 2.01 (n = 30 meiocytes) (one-way ANOVA with Dunnett correction for multiple comparison, both P < 0.001) in *zyp1a/b-1 and zyp1a/b-2* mutants, respectively (Figure 6B and C, Supplemental movie 2). These data suggest that ZYP1 has a dual function in CO formation. On the one hand, ZYP1 assures the formation of at least one CO for each pair of homologs as indicated by the reduced number of chiasmata and enhanced occurrence of univalents in *zyp1 a/b* mutants. On the other hand, it limits the total number of type I COs as indicated by the elevated number of MLH1 foci in *zyp1a/b*.

To understand whether the ZYP1B^Δ661-725^-mediated formation of the SC, which fails to recruit PCH2 for ASY1 removal, would affect the number of type I COs in *zyp1* mutants, we counted the MLH1 foci in *ZYP1BD^661-725^* (*zyp1a/b-1*) plants using our MLH1-GFP reporter system. Interestingly, compared to the wildtype (9.93 ± 1.26, n = 44 meiocytes), we found that the number of MLH1 foci in *ZYP1BD^661-725^* (*zyp1a/b-1*) plants also significantly increased to 12.93 ± 2.05 (n = 25 meiocytes, one-way ANOVA with Dunnett correction for multiple comparison, P < 0.001), and displayed no significant difference from the *zyp1a/b* double mutants (Figure 6B and C, Supplemental movie 3). While we cannot fully exclude the possibility that the precise number of type I COs is strongly dependent on the complete polymerization of ZYP1, this result suggests that the increased formation of type I COs in *zyp1* mutants is very likely not attributed to the absence of ZYP1 *per se*, but the prolonged occupancy of ASY1 on chromosome axes.

## Discussion

Developmental processes usually follow a defined order in space and time. A key aspect is unidirectionality, which prevents cells and tissues from being kept in futile loops. Meiosis is a paradigm for this biological organization principle, which avoids, for instance, that meiocytes are held in repeated cycles of DSB formation and repair. However, how unidirectionality is accomplished in meiosis and other biological processes is often not understood. The dynamic localization of meiotic HORMADs, first present on the chromosome axis at early meiosis and subsequently removed as prophase progresses, is very intriguing in this context and raises the question of how this highly defined order of HORMADs’ localization is achieved. Since this dynamic behavior is well conserved across many sexually-reproducing organisms including species from yeast, mammals, and plants (Rosenberg & Corbett, 2015; Chen *et al*, 2014; Wojtasz *et al*, 2009; Lambing *et al*, 2015; Nonomura *et al*, 2006), it is tempting to speculate that the temporally regulated assembly and disassembly of HORMADs is key to meiosis (Rosenberg & Corbett, 2015; Chen *et al*, 2014; Wojtasz *et al*, 2009; Lambing *et al*, 2015; Nonomura *et al*, 2006). Here, we have revealed an elegant mechanism of how ASY1 assembly and disassembly can be temporally separated, driving first the occupation of the chromosome axis with ASY1 at early prophase I and then its coordinated removal when homologs are synapsed. Moreover, this work has shed light on the question what the biological relevance of this dynamic behavior of HORMADs is.

Previous studies have shown that the loading of meiotic HORMADs on the axis depends on their interacting partners, i.e., the coiled-coil axial core proteins such as ASY3/Red1 in Arabidopsis and yeast, respectively (Smith & Roeder, 1997; Yang *et al*, 2006; Ferdous *et al*, 2012). These axial core proteins are proposed to bind via their closure motif to the HORMA domain of the meiotic HORMADs, thereby recruiting the HORMADs to the chromosome axis (Yang *et al*, 2020b; West *et al*, 2019). However, HORMADs also have a closure motif that is hypothesized to binds to their own HORMA domain, thus leading to a so-called closed conformation and precluding the recruitment of HORMADs to the axis (Yang *et al*, 2020b; West *et al*, 2018; Rosenberg & Corbett, 2015). The AAA+ ATPase PCH2 catalyzes the transition of HORMADs from a closed to an open conformation (Ye *et al*, 2017; West *et al*, 2017). PCH2-mediated conformation change is thought to enable the nuclear targeting and chromosomal assembly of ASY1 in Arabidopsis resembling similar functional interplay between Hop1, PCH2 and the SC in yeast (Yang *et al*, 2020b, 2020a; Balboni *et al*, 2020; Herruzo *et al*, 2021).

Intriguingly, PCH2 also catalyzes the disassembly of ASY1 and we have shown here that the functional switch in PCH2 action is brought about by the installation of the SC, which directly recruits PCH2 via binding to the TF protein ZYP1. In contrast, COMET, the adaptor protein for PCH2, does not require ZYP1 for its localization at the axis and probably only ASY1 itself is responsible for the loading of COMET to the axis (Figure 2C) (Balboni *et al*, 2020). Thus, despite a direct interaction between PCH2 and COMET, it seems that they do not mutually recruit each other to the SC. Instead, a bipartite loading process is at work that brings the two parts of the ASY1-dismantling complex together. It is tempting to speculate that ASY1 bound to the axis is sterically protected from an attack by freely diffusing PCH2 and only the incoming SC causes a conformational change of ASY1 and/or stabilizes the interaction between COMET and PCH2 to allow the efficient removal of ASY1 from the axis. By this mechanism, progression of meiosis is linked with the establishment of a new protein complex providing a means to the unidirectionality of meiosis. Other examples of this biological organization principle are the formation of the DNA replication complexes and the assembly of the spliceosome, for which the next step can only be executed if the preceding protein is correctly loaded and, at the same time, the binding of the following protein often causes the release of the previously bound factor (Siddiqui *et al*, 2013; Matera & Wang, 2014).

Notably, orthologs of PCH2 in different organisms exhibit a similar SC-dependent localization pattern (Joshi *et al*, 2009; Silva & Vader, 2021; Raina & Vader, 2020; Miao *et al*, 2013), and we have shown here that TRIP13, the ortholog of PCH2 in mouse, binds directly to the C-terminus of the murine ZYP1 homolog SYCP1. Therefore, we postulate that a similar mechanism controls SC dependent axial depletion of HORMADs in Arabidopsis and other organisms, such as mammals.

Deleting the binding site of PCH2 in ZYP1 allowed us then to shed light on the question of which aspects of *zyp1* mutants are related to a malfunctioning SC as opposed to secondary consequences, such as the failure to remove ASY1 and subsequent consequences. Moreover, a comparison of *pch2* mutants with the *zyp1* separation-of-function allele allowed us to understand which aspects of *pch2* mutants are related to prolonged presence of ASY1 at the chromosome axis versus additional functions of PCH2 in meiosis.

The finding that the incomplete removal of ASY1 does not necessarily lead to compromised chromosome coalignment/synapsis and chromosome missegregation as seen in *pch2* mutants (Lambing *et al*, 2015; Cuacos *et al*, 2021) indicates that the meiotic defects of *pch2* mutants are not only due to a defective release of ASY1. However, the meiotic defects of *pch2* mutants still appear to be largely related to ASY1 since PCH2 is also required for the efficient nuclear targeting of ASY1 in early prophase and hence promotes meiotic recombination and synapsis (Yang *et al*, 2020a; Cuacos *et al*, 2021).

At the moment, the complexity and consequences of a reduced ASY1 loading together with a compromised removal are difficult to dissect. We cannot exclude the possibility that, in Arabidopsis, PCH2 has further roles in meiosis, such as the involvement in a meiotic checkpoint by monitoring synapsis and meiotic recombination as reported in budding yeast (Ho & Burgess, 2011; San-Segundo & Roeder, 1999; Raina & Vader, 2020; Herruzo *et al*, 2021, 2019). Indeed, it was recently shown that Arabidopsis and likely other plants also have a pachytene checkpoint (Jaeger-Braet *et al*, 2021). However, whether this checkpoint relies on PCH2 needs to be tested in future.

An important aspect of *zyp1* mutants is the increase in type I COs in comparison to the wildtype (France *et al*, 2021; Capilla-Pérez *et al*, 2021). Possibly, removal of ASY1 discharges DSB sites preventing excess DSBs and/or destabilizes the recombination machinery (Vrielynck *et al*, 2021; Sanchez-Moran *et al*, 2007). Support for such a function comes from the recent finding that ASY1 directly interacts with several DSB proteins such as MTOPVIB, PRD2, and PRD3 (Vrielynck *et al*, 2021), and possibly anchors these proteins to the axis. Moreover, loss of *ASY1* shortens the residence time of the recombinase DMC1 on chromosomes at early meiosis (Sanchez-Moran *et al*, 2007), suggesting that ASY1 is likely involved in the stabilization and/or recruitment of (at least some of) the components of the recombination machinery. Therefore, the removal of ASY1 and possibly its homologs in other species represents another means of introducing unidirectionality by terminating meiotic recombination. It is then crucial that the re-association of ASY1 with the axis is prevented possibly by the sequestration of PCH2 to ZYP1 and a fast self-closing of ASY1 present in the nucleoplasm (Yang *et al*, 2020a).

In addition to an increase in CO number, France *et al* (2021) and Capilla-Pérez *et al* (2021) showed that, compared to the wildtype, CO interference, the mechanism that restrains the COs occurring in close proximity, is reduced in *zyp1* mutants. However, it remains unclear how the SC modulates interference. One possibility would be that ZYP1, as it nucleates between the paired axes to form the SC, directly conveys a signal through an unknown mechanism. According to the beam-film model for CO interference, the polymerization of ZYP1 might directly or indirectly be involved in the distribution of mechanical stress/force along the entire chromosome modulating the strength of CO interference (Zhang *et al*, 2014; Wang *et al*, 2015; Kleckner *et al*, 2004). Alternatively, the SC might be a liquid crystal that provides a permissive environment for the diffusion and positive feedback between pro-CO factors. Positive feedback may result in one or a few sites where pro-CO factors reach critical levels for the differentiation of CO-specific intermediates on each homolog pair (Qiao *et al*, 2014; Morgan *et al*, 2021).

Interestingly, ASY1 was previously shown to be essential for mediating CO interference since the COs are very closely located in *asy1* mutants (Lambing *et al*, 2020a; Pochon *et al*, 2022). Thus, both the prolonged presence of ASY1, as seen in the *ZYP1* separation-of-function allele, and the absence of ASY1 from the axes, as in *asy1* mutants, contribute to CO interference. Nonetheless, these two points might not be comparable and mutually exclusive since, compared to the largely normal homologous pairing/coalignment in *zyp1* mutants, the COs are restricted to the synapsed regions of *asy1* mutants at the chromosome ends (Lambing *et al*, 2020a; Ferdous *et al*, 2012; Pochon *et al*, 2022).

Notably, the loss of *ASY1* was found to reduce CO assurance, and the appearance of univalents in *zyp1* mutants suggests that CO assurance is also affected since, in theory, sufficient COs (even more than in the wildtype) formed so that each homologous chromosome pair received at least one CO. The reason for the CO assurance defects of *zyp1* are not clear yet but possibly could go back to the finding that, although chromosomes co-align in *zyp1* mutants, the homologs are not as closely juxtaposed as in the wildtype in pachytene (France *et al*, 2021; Capilla-Pérez *et al*, 2021), presumably destabilizing and/or reducing the resilience of the homolog interaction. Future work, especially using super resolution imaging, may shed light on this aspect and could help to define a distance threshold that allows CO formation. Importantly, our separation-of-function allele of *ZYP1* suggested that the CO assurance defect in *zyp1* mutants is largely due to the lack of synapsis and not the persistence of ASY1 at the chromosome axis. This in turn could also explain why *asy1* mutants, which are asynaptic, CO assurance is also largely compromised.

## Acknowledgements

We thank Attila Toth (Technical University Dresden, Germany), Ihsan Dereli (Max Planck Institute for Multidisciplinary Sciences, Göttingen, Germany) and Mariana Motta as well as Jason Sims (both University of Hamburg, Germany) for the critical reading of the manuscript. We are grateful to Attila Toth for providing murine TRIP13 and SYCP1 cDNAs. C.Y. was supported by funds from the National Natural Science Foundation of China (32170354) and Huazhong Agricultural University. C.Y. received a postdoc-first grant of the Department of Biology of the University of Hamburg. A.S. was supported by funds from the University of Hamburg.

## Author contributions

C.Y. conceived the research and designed the experiments. C.Y. performed most of the experiments. K.S., B.H., and M.B. carried out chromosome spreads, Y.H. performed additional microscopy work, H.T.E. conducted the Y2H analysis of murine proteins, A.S. provided reagents and chemicals, C.Y., H.M., and A.S. analyzed the data. C.Y. and A.S. wrote the manuscript.

---

**Supplemental figure 1.**
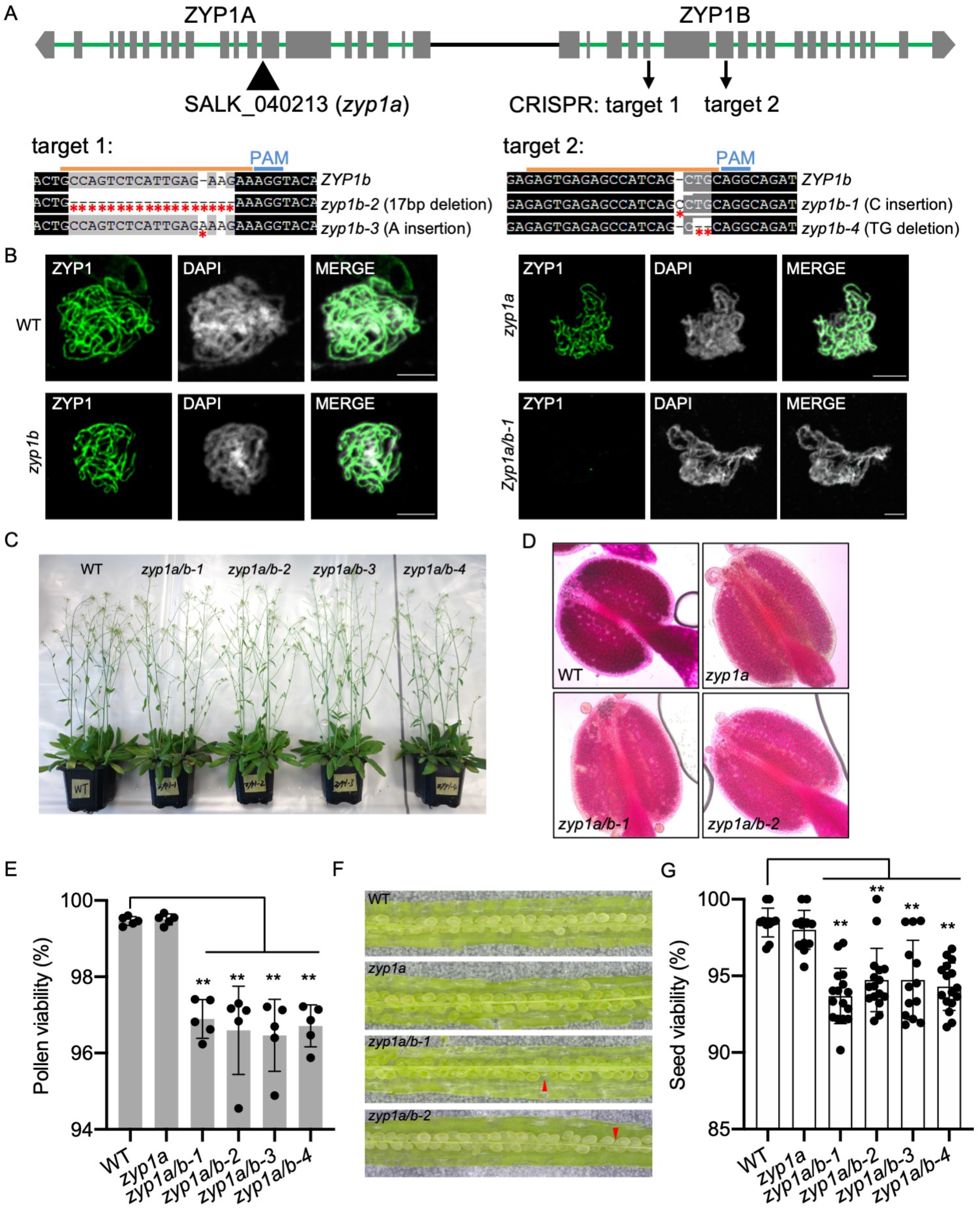
Phenotypical analysis of *zyp1* mutants. (A) Generation and mutation identification of *zyp1a/b* double mutants by CRISPR-Cas9. (B) Immunostaining of ZYP1 in WT, *zyp1a, zyp1b*, and *zyp1a/b-1* mutant plants. Bars: 5 μm. (C) Plants of WT, *zyp1a/b-1, zyp1a/b-2, zyp1a/b-3*, and *zyp1a/b-4* at flowering stage. (D) Pollen staining of WT, *zyp1a, zyp1a/b-1*, and *zyp1a/b-2* mutant plants. Red staining indicates viable pollen and blue indicates dead pollen. (E) Pollen viability of WT, *zyp1a, zyp1a/b-1, zyp1a/b-2, zyp1a/b-3*, and *zyp1a/b-4* mutant plants. Asterisks indicate significant difference (P < 0.01). (F) Seed set of WT, *zyp1a, zyp1a/b-1*, and *zyp1a/b-2* mutant plants. Red arrowheads indicate aborted seeds. (G) Seed viability of WT, *zyp1a, zyp1a/b-1, zyp1a/b-2, zyp1a/b-3*, and *zyp1a/b-4* mutant plants. Asterisks indicate significant difference (P < 0.01).

**Supplemental figure 2.**
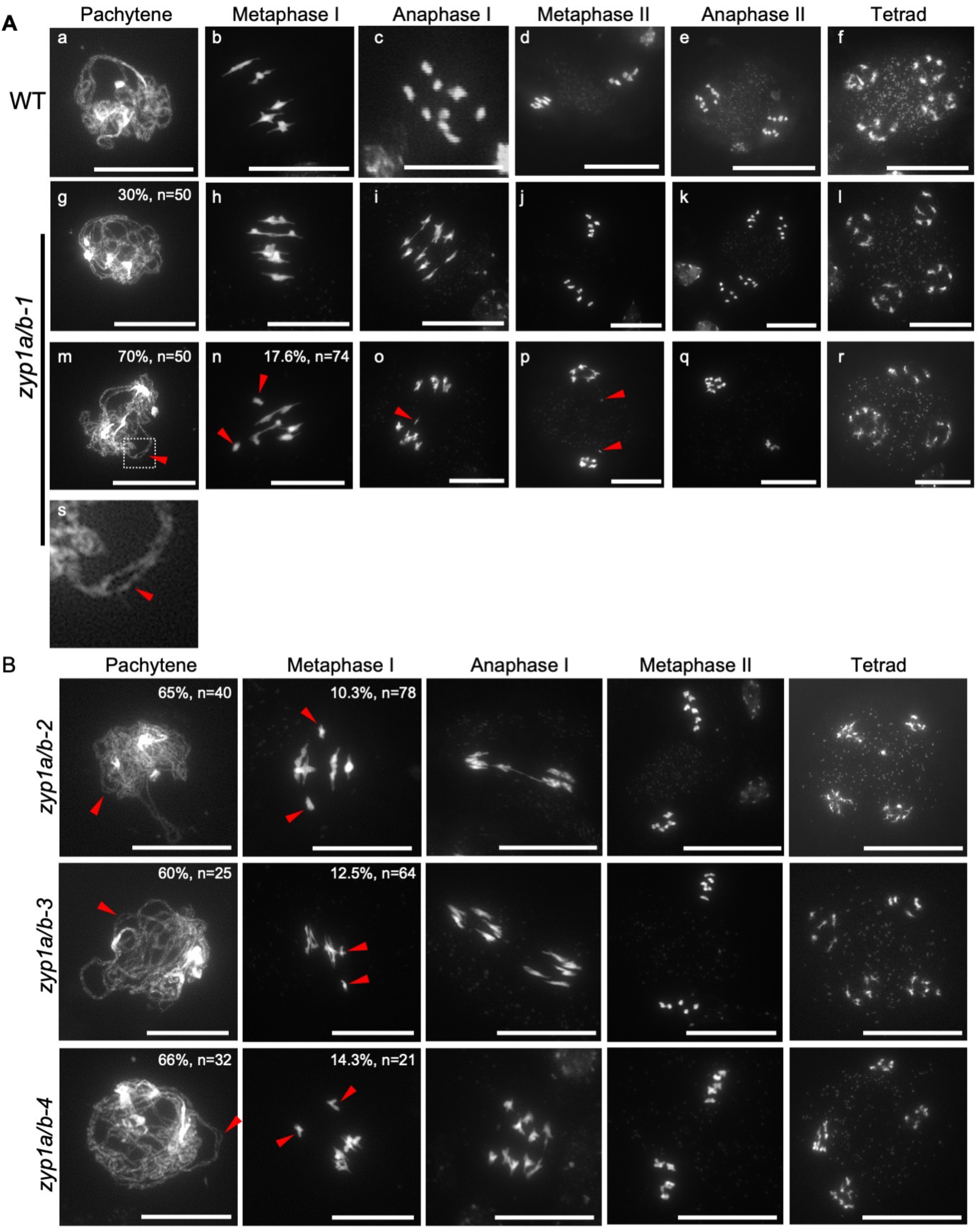
Meiotic chromosome behavior in *zyp1* mutants. (A) Chromosome spreading analysis of male meiosis in WT and *zyp1a/b-1* mutant plants. The pictures of g-l and m-r indicate the normal and abnormal chromosome behaviours in *zyp1a/b-1* mutants, respectively. Red arrowheads indicate the gap between paired chromosomes (m, s), univalents (n), and chromosome fragments (o, p), respectively. Bars: 20 μm. (B) Chromosome spreading analysis of male meiocytes of *zyp1a/b-2, zyp1a/b-3*, and *zyp1a/b-4* mutants. Red arrowheads indicate the gap between paired chromosomes at pachytene or univalent at metaphase I. Bars: 20 μm. The numbers shown in the pictures of pachytene and metaphase I of *zyp1* mutants indicate the number of observed cells showing similar chromosome behaviours and the corresponding percentage.

**Supplemental figure 3.**
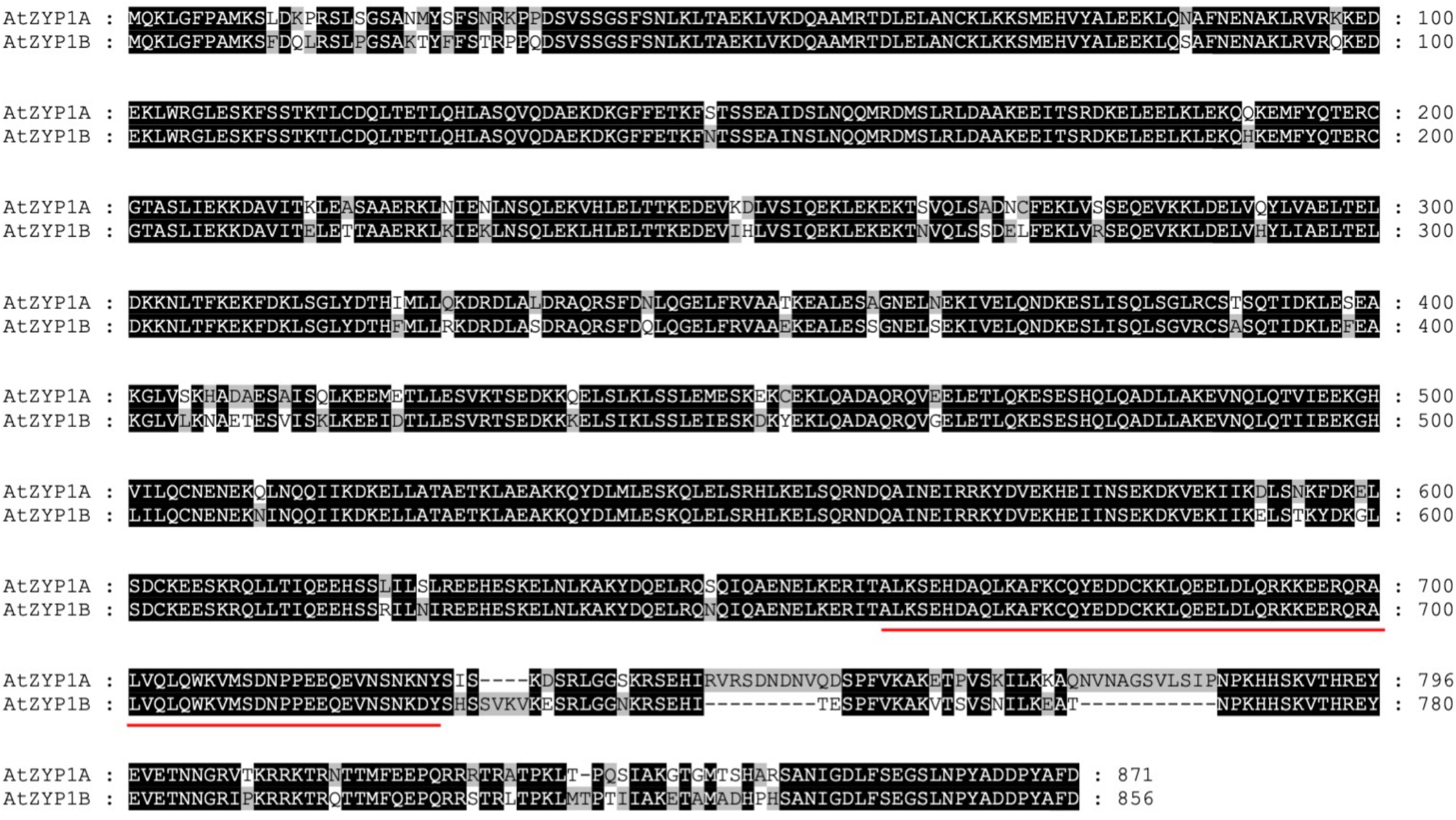
Alignment of ZYP1A and ZYP1B proteins in Arabidopsis. Red lines indicate the amino acids of 661-725 in ZYP1A and ZYP1B.

**Supplemental figure 4.**
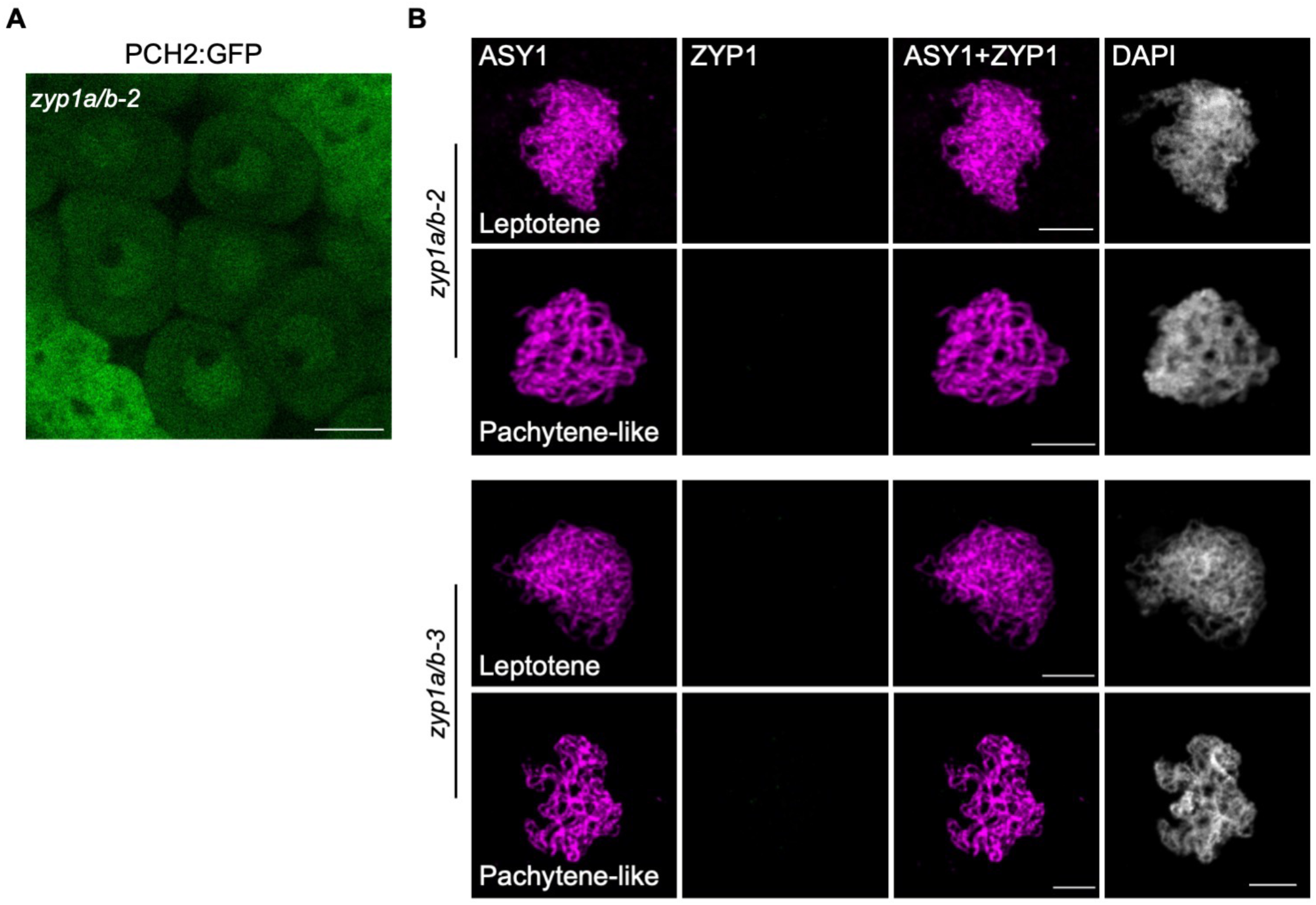
Localization of PCH2 and ASY1 in *zyp1a/b* mutants. (A) Localization of PCH2:GFP in the male meiocytes of the *zyp1a/b-2* mutant at pachytene-like stage. Bar: 10 μm. (B) Immunostaining of ASY1 and ZYP1 in *zyp1a/b-2* and *zyp1a/b-3* mutants. Bars: 5 μm.

**Supplemental figure 5.**
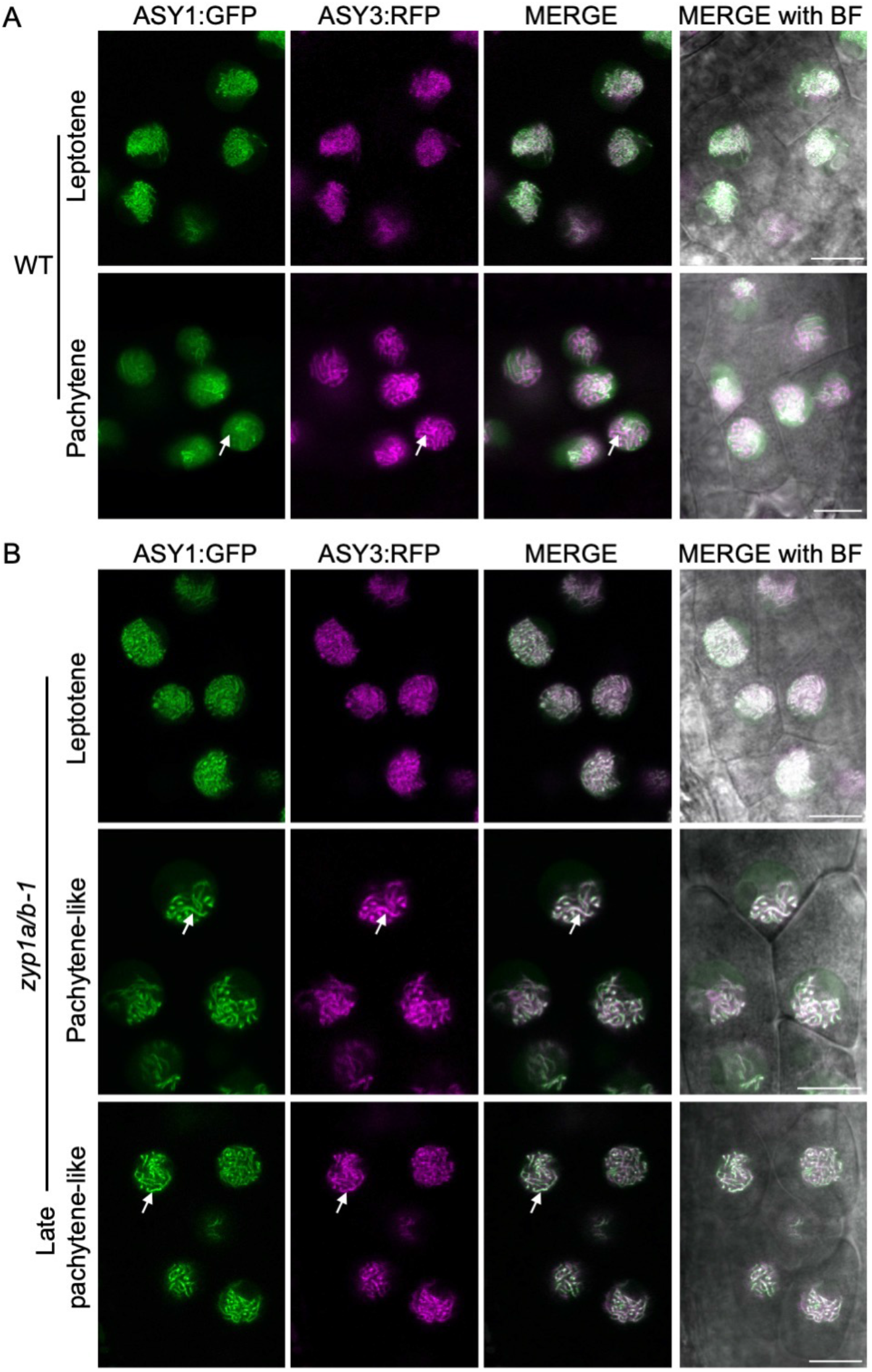
ASY1 removal is compromised in *zyp1* null mutants. (A and B) Localization of ASY1:GFP together with ASY3:RFP in male meiocytes of WT (A) and *zyp1a/b-1(B*) mutant plants at different prophase I stages using confocal laser scanning microscope. BF represents the channel of bright field showing the cell morphology. White arrows indicate chromosome region where ASY1:GFP is largely depleted from the synapsed chromosomes at pachytene (B), and the region where ASY1:GFP proteins show prolonged stay on paired chromosomes at pachytene-like stage. Bars: 10 μm.

**Supplemental figure 6.**
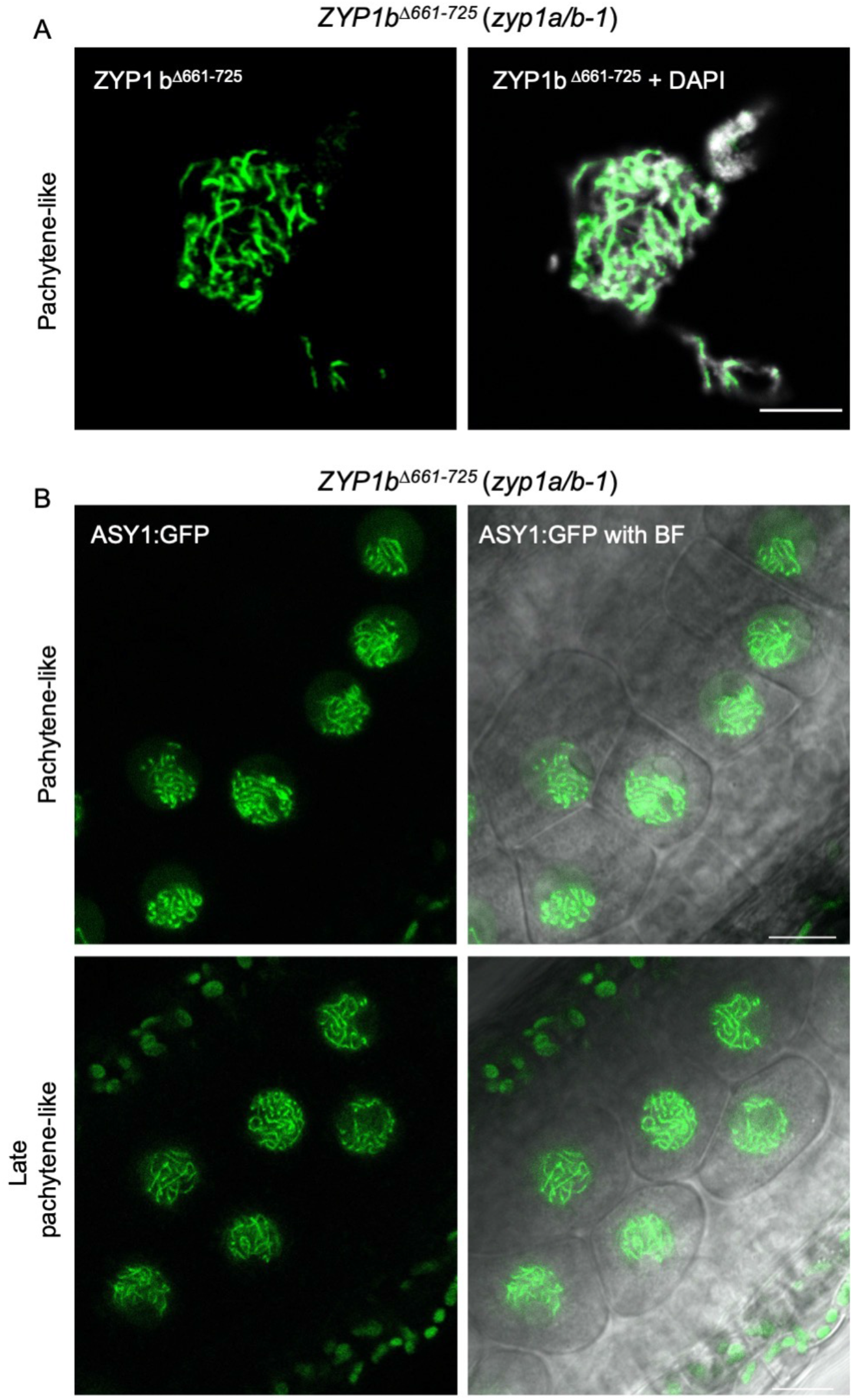
*ZYP1B*^*Δ661*-725^ (*zyp1a/b-1*) shows a largely normal synapsis, but is defective in ASY1 removal. (A) Immunostaining of ZYP1 in the male meiocytes of *ZYP1B^Δ661-725^* (*zyp1a/b-1*) mutant at pachytene-like stage. Bars: 5 μm. (B) Localization of ASY1:GFP in the male meiocytes of *ZYP1B^Δ661-725^* (*zyp1a/b-1*) mutant using confocal laser scanning microscopy. The merge shows the overlay of ASY 1:GFP with the bright field (BF). Bars: 10 μm.

**Supplemental movie 1.** Z-stacks of MLH1:GFP in male meiocytes of wild-type plants at late prophase I.

**Supplemental movie 2.** Z-stacks of MLH1:GFP in male meiocytes of *zyp1a/b-1* mutant plants at late prophase I.

**Supplemental movie 3.** Z-stacks of MLH1:GFP in male meiocytes of *ZYP1B^Δ661-725^ (zyp1a/b-1)* mutant plants at late prophase I.

**Supplemental table 1.**
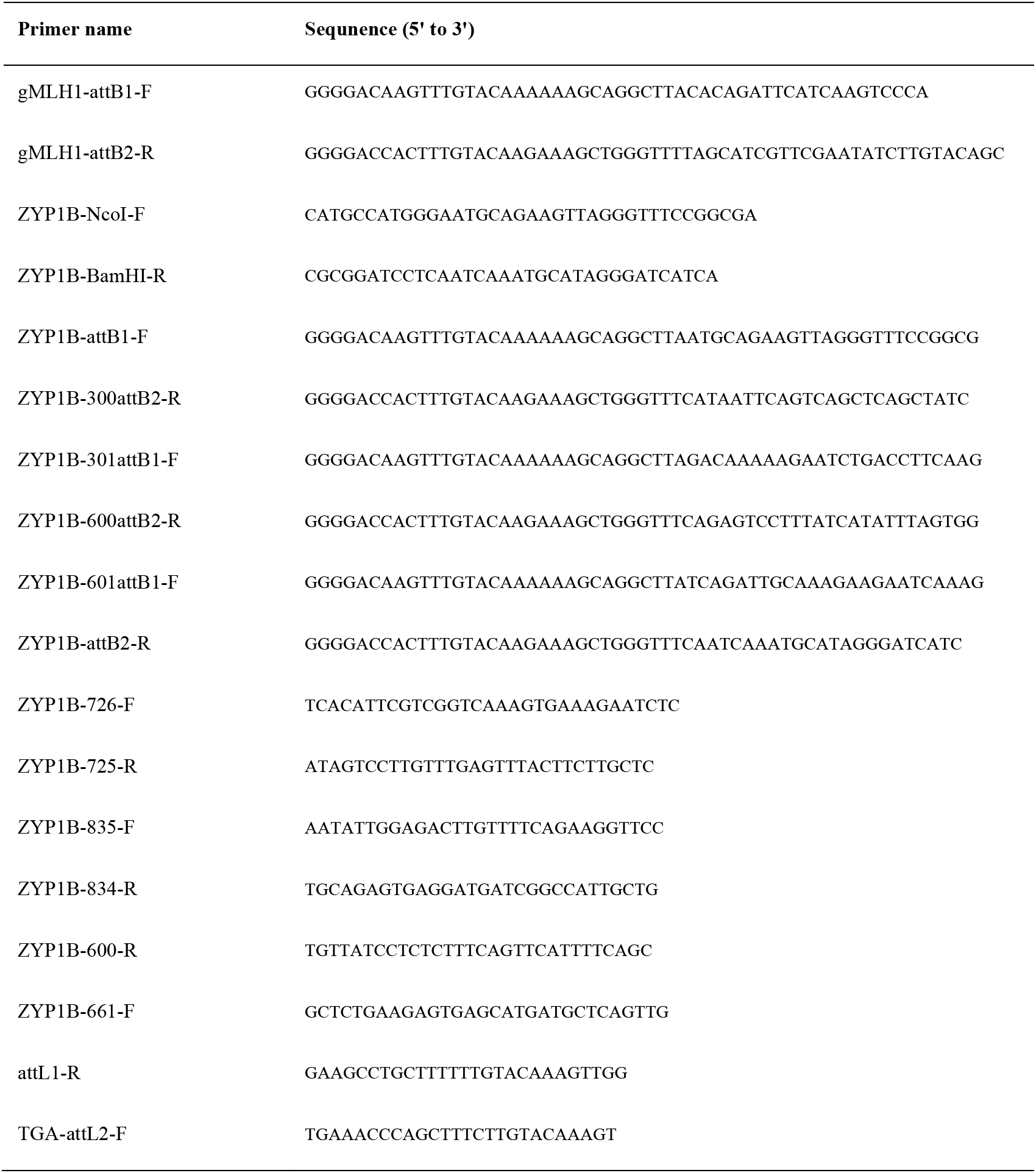
Primers used in this research.

